# Partner-independent fusion gene detection by multiplexed CRISPR/Cas9 enrichment and long-read Nanopore sequencing

**DOI:** 10.1101/807545

**Authors:** Christina Stangl, Sam de Blank, Ivo Renkens, Tamara Verbeek, Jose Espejo Valle-Inclan, Rocio Chamorro González, Anton G. Henssen, Markus J. van Roosmalen, Ronald W. Stam, Emile E. Voest, Wigard P. Kloosterman, Gijs van Haaften, Glen Monroe

## Abstract

Fusion genes are hallmarks of various cancer types and important determinants for diagnosis, prognosis and treatment possibilities. The promiscuity of fusion genes with respect to partner choice and exact breakpoint-positions restricts their detection in the diagnostic setting, even for known and recurrent fusion gene configurations. To accurately identify these gene fusions in an unbiased manner, we developed FUDGE: a FUsion gene Detection assay from Gene Enrichment. FUDGE couples target-selected and strand-specific CRISPR/Cas9 activity for enrichment and detection of fusion gene drivers (e.g. *BRAF, EWSR1, KMT2A*/*MLL*) - without prior knowledge of fusion partner or breakpoint-location - to long-read Nanopore sequencing. FUDGE encompasses a dedicated bioinformatics approach (NanoFG) to detect fusion genes from Nanopore sequencing data. Our strategy is flexible with respect to target choice and enables multiplexed enrichment for simultaneous analysis of several genes in multiple samples in a single sequencing run. We observe on average a 508 fold on-target enrichment and identify fusion breakpoints at nucleotide resolution - all within two days. We demonstrate that FUDGE effectively identifies fusion genes in cancer cell lines, tumor samples and on whole genome amplified DNA irrespective of partner gene or breakpoint-position in 100% of cases. Furthermore, we show that FUDGE is superior to routine diagnostic methods for fusion gene detection. In summary, we have developed a rapid and versatile fusion gene detection assay, providing an unparalleled opportunity for pan-cancer detection of fusion genes in routine diagnostics.

## Introduction

Fusion genes are hallmarks of many human cancers. Recent studies suggest that up to 16% of cancers are driven by a fusion gene^1^. Some cancer types, such as prostate cancer or chronic myeloid leukemia, are characterized by a specific fusion gene (*TMPRSS2-ERG* and *BCR-ABL1* respectively), whereas other cancer types do not show such clear associations ^1,2^. Most fusion genes are highly variable with respect to fusion gene configurations and exact breakpoint-locations. Often, one gene is a recurrent fusion partner (e.g. *KMT2A*/*MLL, ALK*) which exhibits a tissue-specific pattern^3^. However, these genes can fuse to a multitude of partners to obtain their oncogenic potential. One striking example is *KMT2A*, formerly known as MLL, which is a prominent fusion partner in pediatric acute myeloid leukemia (AML) and the predominant fusion partner in acute lymphocytic leukemia (ALL) diagnosed in infants (i.e. children <1 year of age), and has been reported with more than 130 different gene configurations^4,5^.

Whereas fusion detection is pathognomonic for some cancer types, it is a determinant of prognosis or treatment choices in other cancer types^6^,^7^. However, the high levels of variability in fusion gene configurations drastically limits diagnostic detection. Current diagnostic strategies include (break-apart) Fluorescence In Situ Hybridization (FISH) and reverse transcription quantitative polymerase chain reaction (RT-qPCR) assays, depending on the knowledge and breakpoint-variability of the fusion partner^7^. However, these assays are laborious and time-consuming and may not identify the fusion partner. Targeted next generation sequencing (NGS) assays overcome these limitations partially, but are accompanied with a longer turnaround-time, increased costs and bioinformatic challenges. Recent long-read sequencing technologies such as Oxford Nanopore Technology (ONT) sequencing have proven immensely helpful in elucidating structural variation in human genomes^8^. Furthermore, the real-time sequencing capabilities yield abundant opportunities for clinical applications. However, sequencing throughput from one Nanopore flow cell (2-5x genome coverage) is insufficient to elucidate the complete structural variation (SV) landscape of a genome^9^. ONT recently released a Cas9-based protocol for enrichment of specific genomic regions, which utilizes the upstream (5’) and downstream (3’) flanking sequences of the region of interest (ROI), to excise the latter and perform targeted sequencing^10^. Two publications have utilized this method to study methylation and structural variants^10^ as well as genome duplications^11^. With this technique, a median on-target coverage of 165x and 254x was achieved, respectively, offering a unique tool to sequence SVs such as fusion genes. However, this approach requires knowledge of both flanking sequences of the ROI, which again restricts its application to detection of only known fusion gene partner combinations. We here developed FUDGE (FUsion gene Detection assay from Gene Enrichment) as a fusion gene identification strategy to perform targeted enrichment of fusion genes and identify - without prior knowledge - the unknown fusion partner and precise breakpoint by using long-read, real-time ONT sequencing. Furthermore, we created and implemented a complementary bioinformatic tool, NanoFG, to detect fusion genes from long-read Nanopore sequencing data. Utilizing this approach, we are able to achieve an average on-target coverage of 67x - resulting in an average enrichment of 508x - and identify fusion gene partners from various cancer types (e.g. AML, Ewing Sarcoma, Colon) within 48 hours. Additionally, we offer strategies for low-input DNA samples (10 ng) as well as multiplexing of samples and targets to minimize assay costs. Finally, we utilized this method on material in which routine diagnostic procedures were unable to detect the fusion partner, and identified the fusion partner within 2 days.

## Results

### Schematic overview of fusion gene detection assay

We developed FUDGE to specifically enrich for fusion genes in which only one gene partner is known and for which the other fusion gene partner and/or breakpoint is unknown. To achieve this, genomic DNA is dephosphorylated as previously described^10^ and a crRNA flanking the suspected breakpoint region(s) is utilized to target Cas9 to a specific genomic loci where it creates a double-strand DNA break (**Fig. 1A**). The Cas9 protein stays predominantly bound to the PAM-distal side of the cut, therefore masking the phosphorylation side on this end, while exposing phosphorylated DNA on the PAM-proximal side of the cut (**Fig.1B**). This phosphorylated DNA, following dA-tailing, creates a specific contact-point that can be used to anneal the ONT-specific sequencing adaptors - specifically to this region only. To achieve directionality, the crRNAs are designed in a strand-directed manner to specifically direct reads up- or downstream of the crRNA sequence - effectively sequencing into the suspected 5’ or 3’ fusion partner (**Fig.1B, Methods, and Suppl. Fig.1**). Thereafter, the enriched libraries are sequenced on one ONT flow cell. To robustly detect fusion genes from low coverage Nanopore sequencing data, we developed a bioinformatic tool, NanoFG, which reports fusion partners, exact breakpoint-locations, the breakpoint-sequence and primers for validation purposes (**Fig. 1C**).

**Figure 1:**
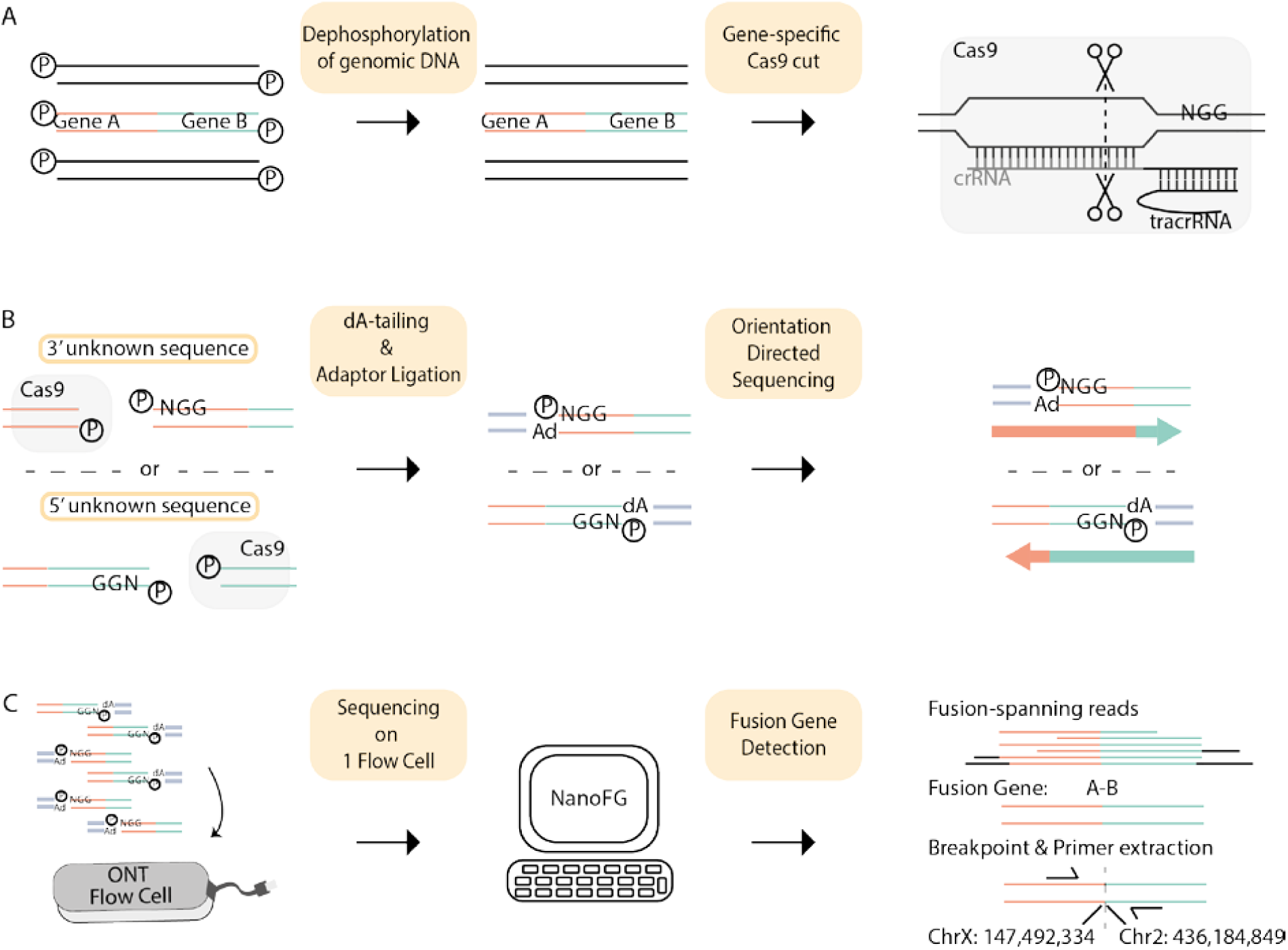
Schematic overview of FUDGE. (**A**) Genomic DNA sample is dephosphorylated and crRNA-guided target-specific double-stranded cuts are introduced through Cas9. (**B**) Phosphorylation-sites are exposed through the double-strand breaks; however, Cas9 remains bound to the PAM-distal side of the cut and blocks phosphorylation of the DNA on this side. DNA-ends are dA-tailed and adaptors are ligated only to phosphorylated DNA-ends proximal to the PAM sequence. Sequencing direction is dictated by the adaptors, towards the unknown sequence. **(C)** Targeted libraries are loaded and sequenced on one ONT flow cell. NanoFG is run on the Nanopore sequencing data, extracts fusion-spanning reads, detects fusion genes and provides exact fusion gene configuration, breakpoint-location, breakpoint-sequence and fusion-spanning primer sequences.

### Genomic enrichment and directed sequencing with single-edge Cas9 targeting

To test the ability of the fusion gene detection assay to generate sufficient enrichment and to direct reads in the desired direction, we applied FUDGE to genomic DNA from a male healthy donor. As a proof-of-principle we designed crRNAs for a panel of recurrent fusion partner genes (*BRAF, EWSR1*, and *SS18*) in a strand-specific manner. We performed two separate library preparations (PP1 and PP2) and targeted two different exons for each of the three genomic loci per library (**Fig. 2A and Suppl. Table 1**). As a positive control, we targeted two genomic loci (*C9orf72* and *FMR1*) for which we previously performed targeted sequencing, and used two crRNAs flanking the ROI and with each targeting one of the two different strands (**Fig. 2A and Suppl. Table 1**). After the sample processing, libraries of PP1 and PP2 were pooled and sequenced a single flow cell. Sequencing resulted in a throughput of 1.665 Gbs which corresponds to a mean genome coverage of 0.5x (**Suppl. Table 1**). For the loci where only one strand of the genome was targeted, on average 89% of the reads sequenced in the anticipated 5’ or 3’ direction (**Fig. 2B-D and Suppl. Fig. 2A-E**). The mean target-locus coverage (10 kb to both sides of the cut-position) was 87x (*BRAF*) (**Fig. 2B**), 96x (*EWSR1*) (**Fig. 2C**), 93x (*SS18*) (**Fig. 2D**), 71x (*C9orf72*) (**Fig. 2E**), and 24x (*FMR1*) (**Fig. 2F**). The average read-length was 9.9kb (**Fig. 2G and Suppl. Table 1**) and on average 116 reads crossed the most common fusion breakpoint-locations (**Fig. 2B-D and Suppl. Table 1**), proving the applicability of this assay to detect fusion genes irrespective of breakpoint-position.

**Figure 2:**
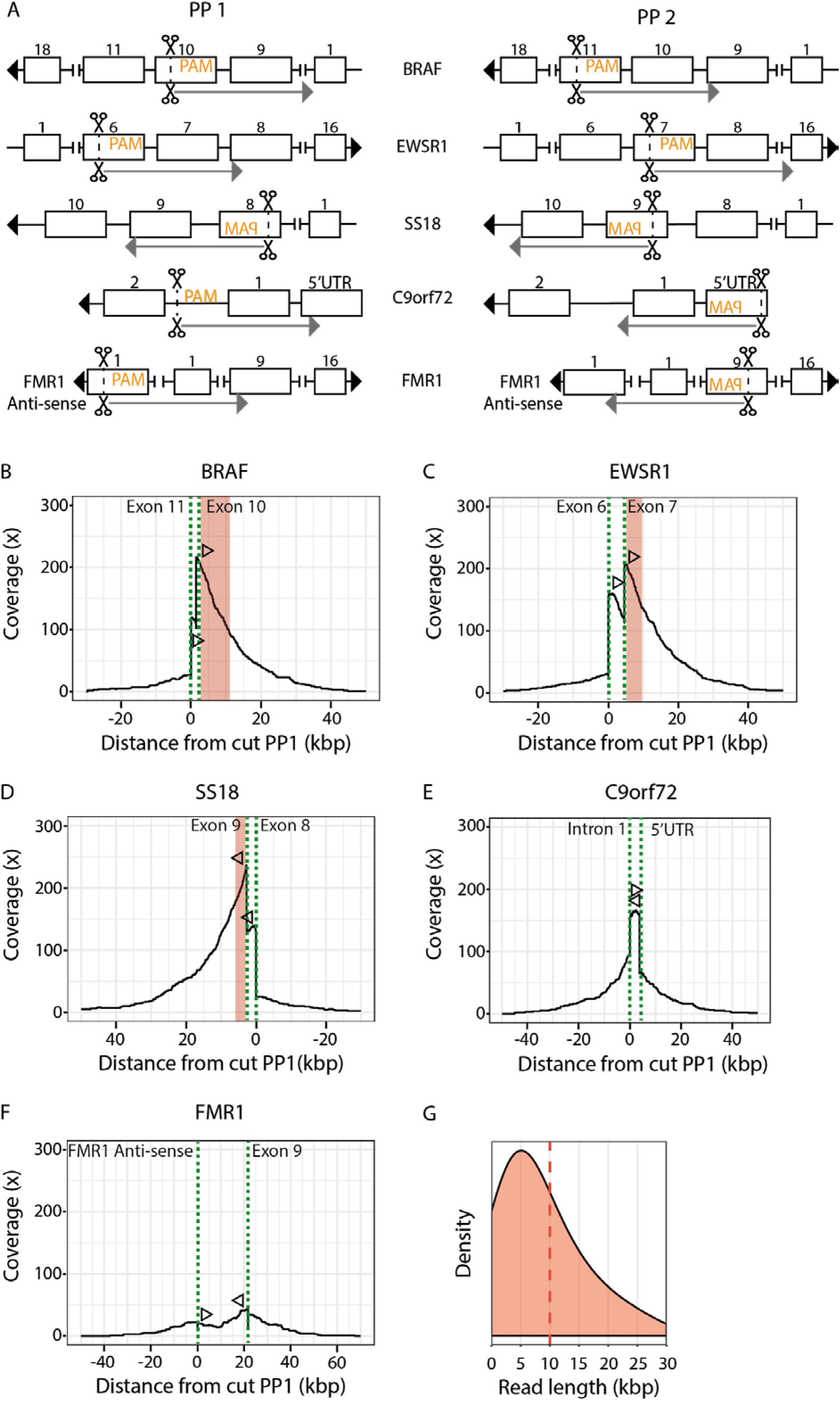
Cas9 enrichment across genomic loci. **(A)** Cas9-introduced cuts targeting different genomic regions for crRNA pool PP1 and PP2. Boxes and numbers represent exons. Cut-positions are shown by scissors, PAM sequence indicates the directionality of crRNA design and arrows show the anticipated sequencing direction. (**B-E**) Coverage plots showing on-target coverage across multiple genomic loci. Dotted lines (green) indicate the crRNA-directed Cas9 cleavage positions and arrows indicate the directionality of reads created from the specific crRNA design. Red areas highlight the most common breakpoint-locations per gene. (**F**) The read-length distribution for the sequencing run. The dashed line indicates the mean read-length.

### Identification of gene fusions in cancer cell lines

To test that FUDGE identifies fusion genes independent of targeted gene or breakpoint-location, we applied this technique to three fusion-positive cancer cell lines in which the fusion configuration was previously identified. The Ewing sarcoma cell lines A4573^12^ and CHP-100^13^ harbour the *EWSR1-FLI1* fusion gene and the synovial sarcoma HS-SYII cell line contains a *SS18-SSX1* fusion^14^. We targeted three loci per sample (*BRAF* Exon 10, *EWS* Exon 7, *SS18* Exon 9) and sequenced the samples on one flow cell each (**Suppl. Table 1**). This produced a mean genome coverage of 0.24x (A4573), 0.14x (CHP-100) and 0.015x (HS-SYII) (**Fig. 3A**). We observed 13x (A4573), 15x (CHP-100) and 4x (HS-SYII) target-locus coverage (10 kb to both sides of the cut-position) and a sharp increase to 81x (A4573), 66x (CHP-100) and 11x (HS-SYII) on-target coverage (cut to breakpoint) due to the achieved directionality (Fig. 3A and Suppl. Fig.1). This relates to an overall on-target fold-enrichment of 342x (A4573), 443x (CHP-100) and 735x (HS-SYII) (**Fig. 3B; Fig 3C-E**). To easily identify fusion-spanning reads from Nanopore data, we developed NanoFG^15^.NanoFG is an amendment to NanoSV^8^ that calls fusion genes from Nanopore sequencing data and reports the exact breakpoint-location, breakpoint-sequence and breakpoint-spanning primers for each gene fusion (**Fig. 1**). NanoFG identified the two *EWSR1-FLI1* fusion genes with 26 (A4573) (**Fig. 3A and 3C**) and 18 (CHP-100) (**Fig. 3A and 3D**) fusion-spanning reads which relates to a fusion-specific enrichment of 109x and 121x, respectively (**Fig. 3B**). The two Ewing sarcoma cell lines harboured the same fusion gene, however, with different breakpoint-locations (**Suppl. Fig. 3**). These differences were readily detected by NanoFG and emphasizes the flexibility of this assay to identify fusions without knowledge of the exact breakpoint-positions. To uncover why NanoFG didn’t identify the *SS18-SSX1* fusion gene, we manually investigated the candidate loci in the IGV Browser^16^. The sequencing of the HS-SYII cell line resulted in very little throughput and on-target coverage (11x) (**Fig. 3A**). As a result, only one fusion-spanning read was produced, which is below the filtering cut-off for fusion-supporting reads set for NanoFG (requirement of minimal two fusion-supporting reads). When adjusting the settings of NanoFG to one supporting read, the *SS18-SSX* fusion was called (**Fig. 3A and 3E**), however, lowering the threshold of fusion-supporting reads requires manual validation if the fusion status is unknown to exclude false-positives. Despite the low-throughput for the HS-SYII cell line, the assay resulted in a 68x fusion-specific fold-enrichment (**Fig. 3B**). This shows the ability of FUDGE to identify fusion genes irrespective of fusion partner or breakpoint-location from low-coverage Nanopore sequencing data.

**Figure 3:**
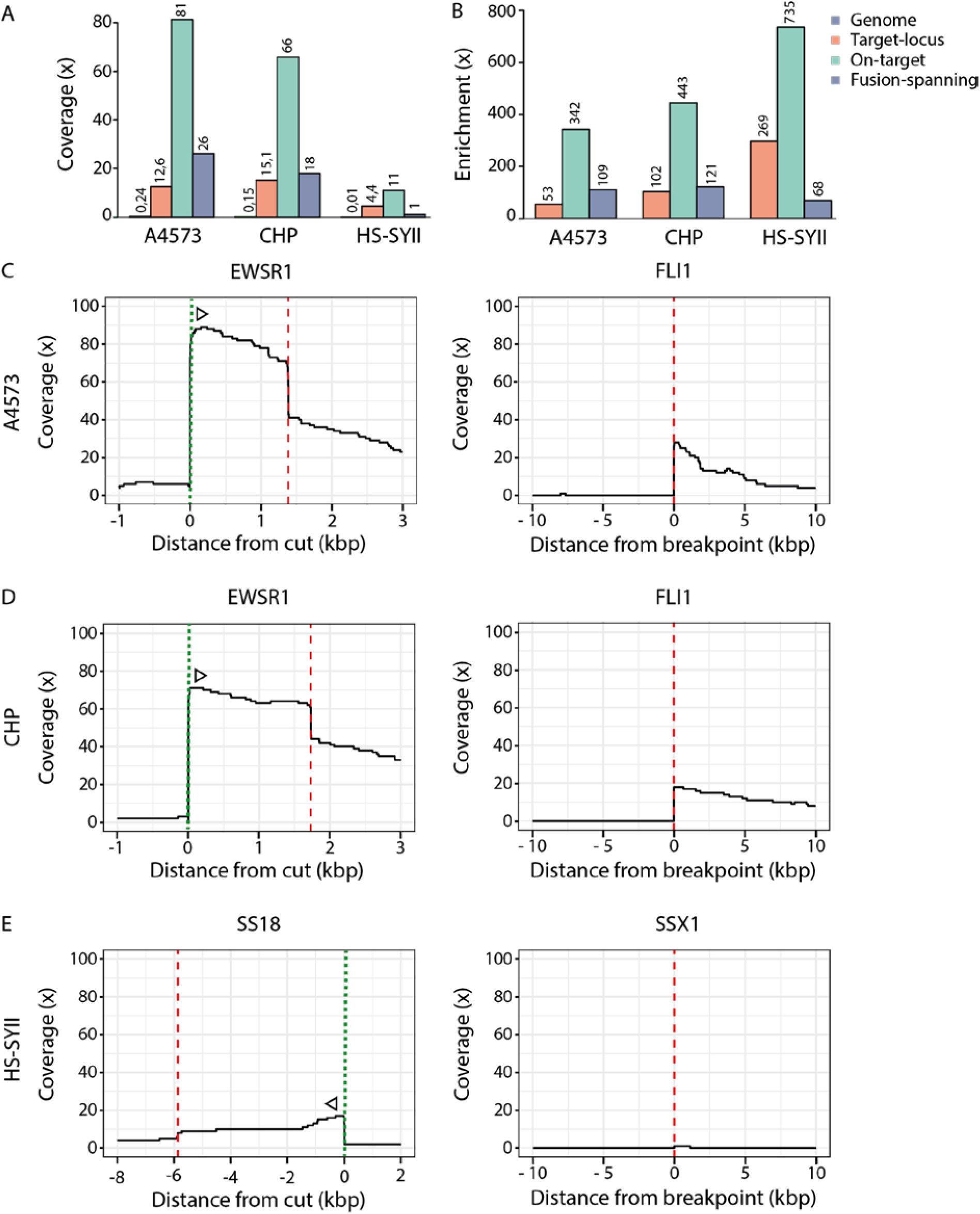
Coverage and enrichment across fusion-positive cancer cell lines. **(A)** Mean coverage and (**B**) enrichment across the genome, target-locus (10 kb to both sides of the cut-position), on-target (cut to breakpoint) and across the fusion junction for the three cell lines A4573, CHP-100 and HS-SYII. (**C**) Coverage plots for the cell line A4573 for the two fusion partners *EWSR1* (targeted) and *FLI1*. (**D**) Coverage plots for the cell line CHP-100 for the two fusion partners *EWSR1* (targeted) and *FLI1*. (**C**) Coverage plots for the cell line HS-SYII for the two fusion partners *SS18* (targeted) and *SSX1*. Dotted lines (green) indicate the crRNA-directed Cas9 cleavage positions and dashed lines (red) indicate breakpoint positions. Arrows indicate the directionality of reads created from the specific crRNA design.

### Detection of fusion genes from tumor material

To validate that FUDGE identifies fusion genes from tumor material and without prior knowledge of the fusion partner, we applied the assay to three tumor samples with (un-)known fusion status. We tested DNA isolated from an Ewing sarcoma (ES1; fusion unknown), a rhabdomyosarcoma (RH; fusion known), and an AML (AML1; one fusion partner known) tumor. Rhabdomyosarcomas are characterized by breaks in the second intron of *FOXO1* (104 kb) which then fuses to either *PAX3* or *PAX7*^*17*^. Due to the large region within *FOXO1* where the break can potentially occur, we chose to target the *PAX3* and *PAX7* genes instead to minimize the number of necessary crRNAs. Here, the most common breakpoint areas span an 18 kb and 32 kb region, respectively. Therefore, we designed sequential crRNAs to span the potential breakpoint regions of both genes (**Suppl. Table. 1**). For the AML sample, diagnostic efforts identified a *KMT2A* fusion through break-apart FISH; however, the fusion partner could not be identified. The KMT2A gene is a frequent fusion partner in AML and ALL and shows two major breakpoint clusters ^4^ for both of which we designed crRNAs (**Suppl. Table 1**). We sequenced each tumor sample on a single flow cell and identified a *EWSR1-FLI1* (ES1) fusion (**Suppl. Table. 1 and Suppl. Fig. 3 and 4**), a reciprocal *FOXO1-PAX3* (RH-1) and *PAX3-FOXO1* (RH-2) fusion (**Fig. 4A**), and a *KMT2A-MLLT6* (AML1) fusion (**Fig. 4B**) with 7, 31, 8 and 25 fusion-spanning reads, respectively (**Fig. 4C**). The reciprocal *FOXO1-PAX3* fusion was validated by breakpoint PCR (**Suppl. Fig. 5A**). On-target enrichment was 498x (ES1), 927x (RH) and 909x (AML1) and the fusion-specific enrichment was 237x (ES1), 150x (RH-1), 65x (RH-2) and 124x (AML1) (**Fig. 4D**). This demonstrates the ability of FUDGE to detect known, unknown and reciprocal fusion genes from patient samples. Furthermore, we performed a retrospective time-course experiment to identify the necessary sequencing time to detect fusion-spanning reads (**Fig. 4E**). On-average, it took 3 hours of sequencing time to identify two fusion-spanning reads, highlighting the speed of our approach.

**Figure 4:**
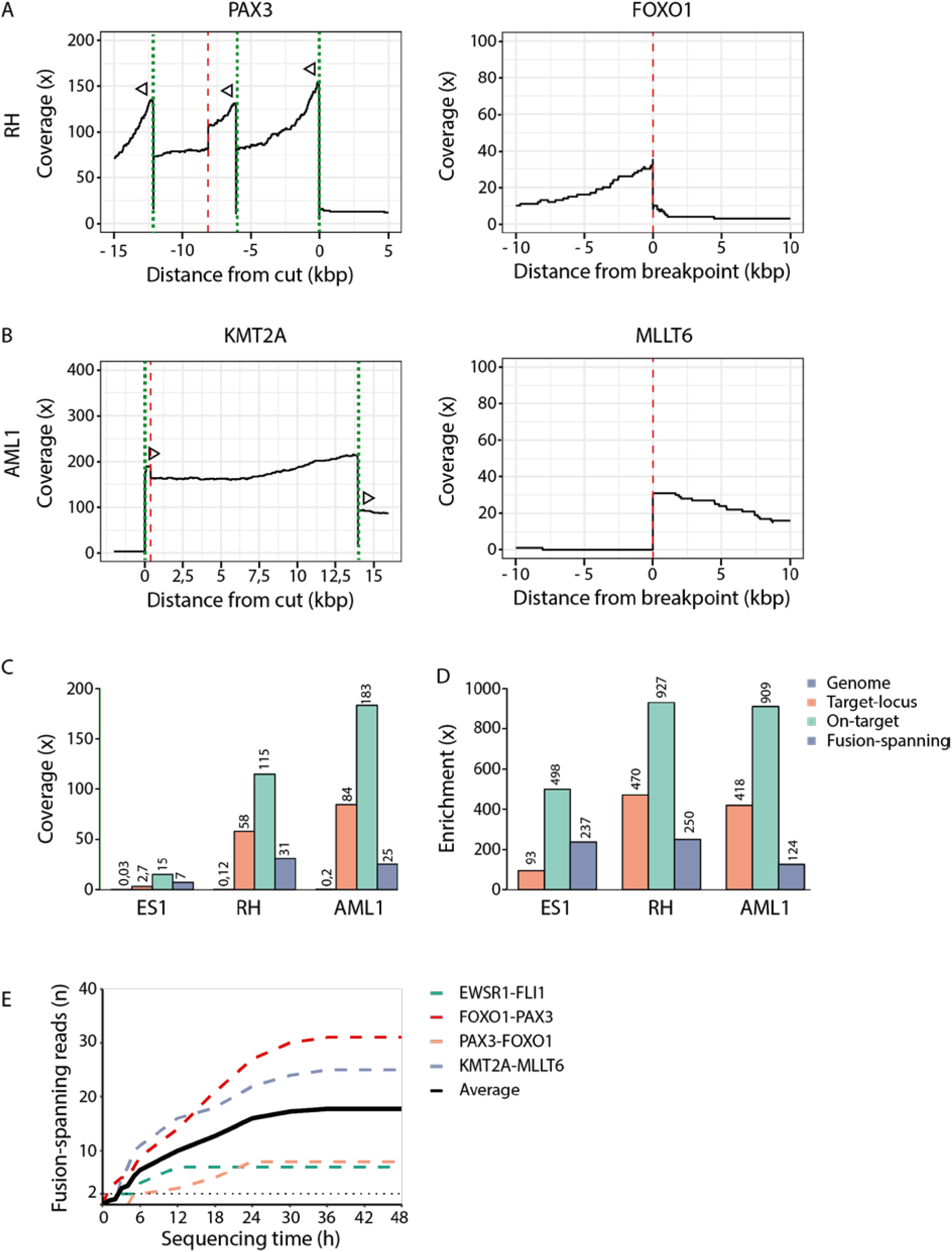
Fusion gene coverage and enrichment from tumor samples. **(A)** Coverage plots for the RH tumor sample for the two fusion partners *FOXO1* and *PAX3. PAX3* was targeted with three sequential guides to span the 18kb possible breakpoint region. **(B)** Coverage plots for the AML1 tumor sample for the two fusion partners *KMT2A* (targeted) and *MLLT6*. (**C**) Mean coverage and (**D**) mean enrichment across the genome, the target-locus (10 kb to both sides of the cut-position), on-target (from cut to breakpoint) and across the fusion junction for the tumor samples ES1, RH and AML1. (**E**) Time-course experiment on sequencing time to identify fusion-spanning reads.

**Figure 5:**
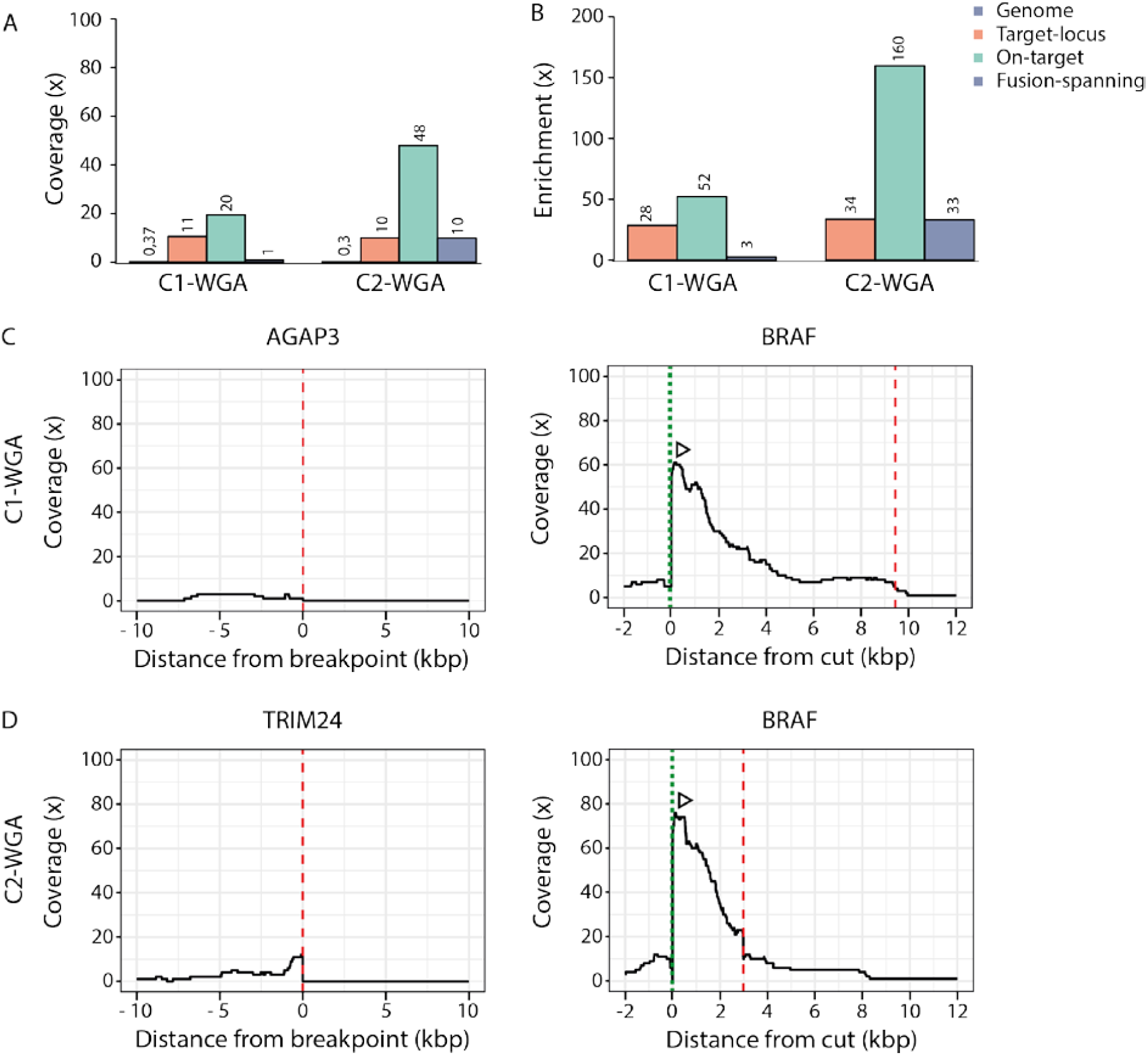
Fusion gene detection from WGA tumour DNA. **(A)** Mean coverage and (**B**) enrichment across the genome, target-locus (10 kb to both sides of the cut-position), on-target (from cut to breakpoint) and across the fusion junction for the whole genome amplified tumor material C1 and C2. (**C**) Coverage plots for the whole genome amplified tumor material C1 for the two fusion partners *AGAP3* and *BRAF* (targeted). (**D**) Coverage plots for the whole genome amplified tumor material C2 for the two fusion partners *TRIM24* and *BRAF* (targeted). Dotted lines (green) indicate the crRNA-directed Cas9 cleavage positions and dashed lines (red) indicate breakpoint positions. Arrows indicate the directionality of reads created from the specific crRNA design.

### Fusion gene detection from low-input tumor material

The amount of available tumour material is often a limiting factor for genomic analysis. To circumvent this problem, we tested if our pipeline was compatible with whole genome amplified (WGA) material. Therefore, we sequenced two colon cancer samples (C1 and C2), known to harbor BRAF fusions (*AGAP3-BRAF* and *TRIM24-BRAF*, respectively)^18^. We performed WGA on 10 ng starting material and subjected 1 ug of WGA-DNA to the enrichment protocol. Genome coverage (**Fig. 5A**) and read-length were comparable to previous experiments (**Suppl. Table 1**). NanoFG detected the the *AGAP3-BRAF* fusion with one fusion-spanning read (**Fig. 5A-C**) and the *TRIM24-BRAF* fusion with 10 fusion-spanning reads (**Fig. 5A-B, 5D**). Of note, WGA introduced unwanted structural variation leading to an increased number fusion gene predictions. Fusion genes identified by NanoFG which were not targeted within our assay are very likely to be false-positives. To validate the two *BRAF* fusion genes, we utilized the exact breakpoint-locations provided by NanoFG and performed breakpoint-spanning PCR on the non-amplified tumor DNA (**Suppl. Fig. 5B**). This demonstrates the power of long-read sequencing to accurately identify structural variants from WGA material. Additionally, for the *BRAF* fusions, the breakpoint junction locations were 6.5 kb apart (**Fig. 5C-D and Suppl. Fig. 3**), highlighting the unbiased performance of our assay. Hence, we show the applicability of our protocol to identify fusion genes from tumor biopsies, even with very limited input material.

### Multiplexing of fusion-positive cell lines

Parallel identification and cost-reduction are key for diagnostic approaches. Therefore, we tested the feasibility to multiplex samples in one sequencing run. We obtained DNA from four *KMT2A*-fusion positive cell lines (ALLPO, KOPN8, ML2 and Monomac-1) with different fusion partners (*MLLT1, MLLT2, MLLT3* and *MLLT4*). We used two crRNAs targeting both breakpoint clusters (**Suppl. Table. 1)** and produced separate libraries for each sample (**Fig. 6A)**. The targeted libraries were pooled pre-sequencing without barcoding and run on a single flow cell. This multiplexing approach resulted in a genome coverage of 0.57x and average read-length of 9.2kb (**Suppl. Table 1**). NanoFG identified the four different fusion partners (**Suppl. Fig. 6A)** and 6 different breakpoint-locations **(Fig. 6B)** Interestingly, two *KMT2A*-fusions (*MLLT2* and *MLLT3*) appeared to be reciprocal (**Suppl. Fig. 6A and Suppl. Fig. 6B**). The breakpoints within *KMT2A* spanned a region of 6 kb, and we identified breakpoints for reciprocal fusions to be location-independent (**Fig. 6C**). We utilized the breakpoint-spanning primers and tested all samples for the occurrence of all fusion genes (**Fig. 6A**). This approach easily deconvoluted the sample-of-origin of each fusion, therefore validating this multiplexing approach (**Suppl. Fig. 7A**). Of note, the Monomac-1 cell line (KMT2A-MLLT3) also exhibited a positive result for the KMT2A-MLLT1 fusion. This could be traced back to a contamination in the cultured cell line, highlighting the sensitivity of this assay to detect subclonal events. Furthermore, from the coverage plot we observed 26 reads within the *MLLT4* fusion partner (**Suppl. Fig. 6A**) which were not explained by any of the NanoFG detected fusions. Upon manual investigation in the IGV browser, we identified one fusion, *KMT2A-MLLT4*, that had a more complex rearrangement which was not called by NanoFG (**Suppl. Fig. 7B**). In this case, a small 30 bp region of *KMT2A* was deleted, followed by a 185 bp inversion and the ultimate fusion to *MLLT4*. We again designed breakpoint-spanning primers and additionally performed Sanger-sequencing on the amplicons and validated the occurrence and structure of the complex rearrangement (**Suppl. Fig. 7B**). As a result, with the use of only one Nanopore flow cell, we identified seven fusion genes from four samples with a collective on-target enrichment of 349x resulting in an average of 18 fusion-spanning reads (Fig. 6D). This shows the ability of our approach to multiplex samples with different fusion genes and breakpoint-positions and pinpoint the sample-of-origin by a simple PCR assay.

**Figure 6:**
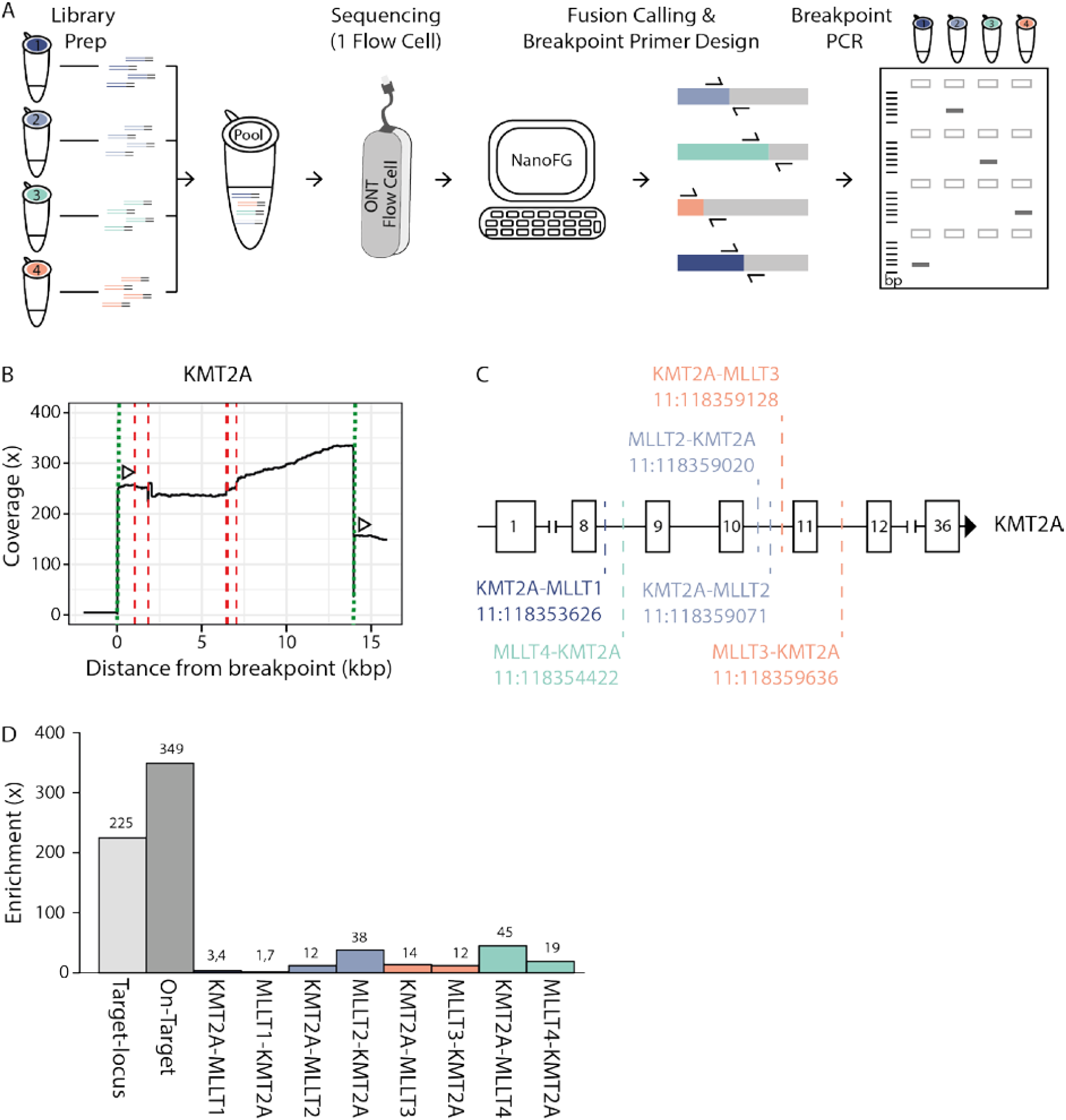
Multiplexing of fusion-positive samples with varying breakpoints and fusion partners. (**A**) Schematic overview of the multiplexing approach. Samples are prepared and subjected to the Cas9-enrichment individually and pooled equally pre-sequencing. The library-pool is sequenced on a single ONT Flow Cell and NanoFG detected fusion genes and designed fusion-specific breakpoint primers. Original samples are subjected to breakpoint PCR to identify the sample-of-origin for each fusion gene. (**B**) Coverage plots for *KMT2A* (targeted). Dotted lines (green) indicate the crRNA-directed Cas9 cleavage positions and dashed lines (red) indicate breakpoint positions. Arrows indicate the directionality of reads created from the specific crRNA design. (**C**) Breakpoint locations within the KMT2A gene for the different identified fusion genes. Breaks cluster between Exon 8 and 12 and reciprocal fusion genes are highlighted in the same color. (**D**) Mean enrichment across the target-locus (10kb to both sides of the cut-position), on-target (from cut to breakpoint) and across the fusion junctions.

## Discussion

Fusion genes are detrimental determinants for diagnosis, prognosis and treatment opportunities for various cancer types^19^. However, fusion gene detection by diagnostic approaches is limited to highly recurrent fusion gene configurations. We here developed FUDGE, a fusion gene detection assay from gene enrichment coupled to Nanopore sequencing, which enables rapid partner- and breakpoint-location independant fusion gene detection within 48 hours.

Common diagnostic approaches for fusion gene detection range from targeted (qPCR) to semi-targeted (FISH) and unbiased (Whole Genome Sequencing (WGS); RNA-Seq) solutions^7,19^. All offer some benefit but are limited in accuracy, resolution, or turnaround-time. qPCR assays offer accurate information on the fusion partners present and the exact breakpoint location; however, due to the large variety of possible fusion partners and breakpoint-positions, only highly recurrent events can be investigated. FISH offers detection of one fusion partner, but is restricted to one gene per test and does not provide information on the fusion partner. WGS and RNA-Seq potentially detects all fusion genes present in a given sample; however, both are hampered by a high turn-around time and WGS can result in high false-positive rates. A combination of these techniques can be utilized for more specific characterization of fusion genes (i.e. FISH followed by qPCR), but at the cost of both time and money.

With FUDGE we offer fast and unbiased fusion gene detection. We successfully identified fusion genes in 100% of the investigated samples independent of cancer type or fusion gene configuration and/or breakpoint-positions. We targeted five recurrent fusion partners and identified 16 unique fusion gene configurations, highlighting the complexity of fusion gene biology. In one case, *KMT2A* was identified as a fusion partner by break-apart FISH through diagnostic efforts; however, the fusion partner was undetectable. We applied FUDGE to the sample and identified *MLLT6* as the fusion partner within 2 days (provided the crRNA was already designed and in-house). Furthermore, FUDGE also detects reciprocal fusion events without additional efforts. In the case of two *BRAF* fusion-positive samples, the breakpoint locations were > 6 kb apart from each other. Conventional methods such as qPCR would have not sufficed to span this large region of possible breakpoint-positions. We integrated an adaptation to the protocol to design sequential guides, offering the opportunity to span large regions of possible breakpoint-locations. For the *FOXO1-PAX3* fusion, we spanned a >20kb region and identified the breakpoint 7,5 kb from the first targeted sequence, highlighting the versatility of FUDGE.

With our assay, fusion detection is possible within 48 hours. Rapid identification of fusion genes is essential for tumor types where fusion genes are pathognomonic such as Ewing sarcoma or synovial sarcoma^19,20^. Hence, early detection allows for early definitive diagnosis and treatment initiation. Furthermore, occurence of a specific fusion gene configuration can be a determinant of prognosis^21^. FUDGE identified all fusion gene configurations within 48 hours, allowing ultrafast diagnosis and treatment initiation. Additionally we show that 3 hours of sequencing are sufficient to identify two fusion-spanning reads, offering the opportunity to reduce the assay time for urgent cases to less than a day.

Intratumoral heterogeneity and tumor purity are likely to influence the lower detection limits of our assay. We set a cut-off of at least two fusion-spanning reads to reliably detect a fusion gene. For the samples HS-SYII and C1, sequencing throughput was very low, resulting in a low on-target coverage. Furthermore, the fusion breakpoints were approximately 6 kb and 9 kb, respectively, from the targeted region, lowering the amount of reads in the breakpoint area. Here, the fusions were only detected with one fusion-spanning read each, requiring the manual validation of the fusion gene by breakpoint PCR. However, with incorporating a multi-crRNA approach and increased efforts from ONT to improve sequencing throughput, FUDGE is expected to improve. Additionally, the latter would allow for higher capacities to multiplex samples, reducing costs of the assay further.

Until now, we focused our assay on five recurrent fusion genes, however, expanding the assay to any gene of interest is possible. Furthermore, rapid detection of the exact breakpoint-positions opens the doors to immediately trace fusion molecules within ctDNA from liquid biopsies and monitor treatment responses and minimal residual disease.

In conclusion, FUDGE identifies fusion genes irrespective of fusion partner or breakpoint-location from low-coverage Nanopore sequencing. With its requirement for only very little amount of tumour material, its ability to multiplex targets as well as samples and its rapid nature, FUDGE overcomes various limitations of current diagnostic assays. Therefore, FUDGE permits initiation of appropriate therapies and options for blood-based minimal residual disease testing within due time after patient presentation.

## Material & Methods

### Cell Lines and Culture

Ewing sarcoma cell lines (A4573, CHP-100) and synovial sarcoma cell line (HS-SYII) were cultured in 5% CO2 in a humidified atmosphere at 37 °C in Dulbecco’s modified medium (DMEM) (Thermo Fisher) supplemented with 10 % fetal bovine serum (FBS) and antibiotics (100 U / ml penicillin and 100 μg / ml streptomycin). The absence of *Mycoplasma sp.* contamination was determined with a Lonza MycoAlert system.

ALL cell lines ALL-PO and KOPN8 and AML cell lines ML2 and Monomac-1 were maintained as suspension cultures in RPMI-1640 medium (Invitrogen), supplemented with 10% or 20% fetal calf serum (FCS) and antibiotics.

### Patient material

The healthy donor (PP) provided oral informed consent. The patients ES1 and RH had been registered and treated according to German trial protocols of the German Society of Pediatric Oncology and Hematology (GPOH). This study was conducted in accordance with the Declaration of Helsinki and Good Clinical Practice, and informed consent was obtained from all patients or their guardians. Collection and use of patient specimen was approved by the institutional review boards of Charité Universitätsmedizin Berlin. Specimen, clinical data were archived and made available by Charité-Universitätsmedizin Berlin.

C1 and C2 were previously sequenced ^18^ and were kindly provided by Prof Ijzermans, Dept of Surgery, Erasmus Medical Center Rotterdam, The Netherlands.

AML1 was a kind gift from Prof. dr. C.M. Zwaan, Erasmus Medical Center – Sophia Children’s Hospital, Rotterdam, The Netherlands / Princess Maxima Center for Pediatric Oncology, Utrecht, The Netherlands. Informed consent is given by the patient or his/her parents or legal guardians, and all is performed in line with the declaration of Helsinki, and the Erasmus MC – Sophia Children’s Hospital approved the experiments.

### DNA-Isolations

Genomic DNA from cultured cells (A4573, CHP-100 and HS-SYII) and tissue (ES1 and RH) was extracted by using the column-based NucleoSpin® Tissue DNA extraction kit (Macherey-Nagel) following manufacturer’s instructions. Sample quality control was performed using a 4200 TapeStation System (Agilent), and DNA content was measured with a Qubit 3.0 Fluorometer (Thermo Fisher).

Genomic DNA from the ALL cell lines (ALLPO and KOPN8), AML cell lines (ML2 and Monomac-1) and AML patient (AML1) was isolated by using the column-based Qiagen DNeasy Blood & Tissue DNA extraction kit (Qiagen) following the manufacturer’s instructions and DNA content was measured with a Qubit 2.0 Fluorometer (Thermo Fisher).

### WGA

For whole genome amplification (WGA), 10 ng starting material was amplified with the repli-g mini kit (Qiagen) according to the manufacturer’s protocol.

### crRNA design

Each potential gene fusion constituted a known fusion partner to be targeted with this enrichment technique, and an (un)known partner to be identified following subsequent sequencing. The known target fusion partners were designated as a 5’ or 3’ fusion partner, dependent upon known literature. Furthermore, the most common breakpoint locations were extracted from a literature search and the most distal breakpoint locations were noted as extreme borders of the targeted area. If the unknown fusion partner was the 5’ partner, crRNAs were designed as the sequence present on the minus strand of the gene (5’->3’) until the PAM sequence. If the unknown fusion partner was the 3’ partner, crRNAs were designed as the sequence present on the plus strand of the gene (5’->3’) until the PAM sequence (**Suppl. Fig. 1**). **1)** Custom Alt-R®□ crRNAs were designed with the Integrated DNA Technologies (IDT) custom gRNA design tool and chosen with maximum on-target and lowest off-target scores (IDT).

### Cas9-Enrichment and Nanopore Sequencing

Cas9 enrichment was adapted from the ONT Cas9 enrichment protocol^10^. In brief, approximately 1 ug of genomic DNA or WGA-DNA (See Suppl. Table 1) was dephosphorylated with Quick calf intestinal phosphatase (NEB) and CutSmart Buffer (NEB) for 10 minutes at 37 °C and inactivated for 2 minutes at 80 °C. crRNAs were resuspended in TE pH7.5 to 100 uM. For simultaneous targeting of multiple loci, crRNAs were pooled equimolarly to 100 uM. Ribonucleoprotein complexes (RNPs) were prepared by mixing 100 uM equimolarialy pooled crRNA pools with 100 uM tracrRNA (IDT) and duplex buffer (IDT), incubated for 5 minutes at 95 °C and thereafter cooled to room temperature. 10 uM RNPs were mixed with 62 uM HiFiCas9 (IDT) and 1x CutSMart buffer (NEB) and incubated at RT for 15 minutes produce Cas9 RNPs. Dephosphorylated DNA sample and Cas9 RNPs were mixed with 10mM dATP and Taq polymerase (NEB) at 37 °C for 15 minutes and 72 °C for 5 minutes to facilitate cutting of the genomic DNA and dA-tailing. Adaptor ligation mix was prepared by mixing Ligation Buffer (SQK-LSK109, ONT), Next Quick T4 DNA Ligase (NEB) and Adaptor Mix (SQK-LSK109, ONT). The mix was carefully applied to the processed DNA sample without vortexing and incubated at room temperature for 25 minutes. DNA was washed and bound to beads by adding TE pH8.0 and 0.3 x volume AMPure XP beads (Agencourt) and incubated for 10 minutes at room temperature. Fragments below 3 kb were washed away by washing the bead-bound solution twice with Long Fragment Buffer (SQK-LSK109, ONT). Enriched library was released from the beads with Elution Buffer (SQK-LSK109, ONT). Enriched library concentration was measured with the a Qubit Fluorometer 3.0 (Thermo Fisher). The library from one tumour sample was loaded onto one Flow Cell (R 9.4, ONT) according to the manufacturer’s protocol. Sequencing was performed on a GridION X5 instrument (ONT) and basecalling was performed by Guppy (ONT).

### NanoFG

NanoFG can be found in https://github.com/SdeBlank/NanoFG

#### Mapping and SV detection

Reads were mapped to the human reference genome version GRCHh37 by using minimap2 (v. 2.6)^22^ with parameters: ‘-x map-ont -a’. The produced SAM file was compressed to bam format and indexed with samtools (v. 1.7)^23^. Next, structural variations were detected from the bam file. The user can choose either NanoSV (v. 1.2.4)^8^ with default parameters: ‘min_mapq=12, depth_support=False, mapq_flag=48’ or Sniffles^23^ with default parameters: ‘-s 2 -n -1 --genotype’ to detect SVs. We here used NanoSV for all experiments (except multiplexing). For the samples C1 and HS-SYII, additional parameters: ‘cluster_count=1’ were used for NanoSV due to the low number of reads spanning the fusion. For the multiplexing experiment, the fraction of reads supporting the fusions was below the allele frequency cut-off in NanoSV. Therefore, the default Sniffles settings were used to detect 6 fusions. By default, all SVs that do not pass the built-in NanoSV or Sniffles filters are removed. Additionally, all insertions are also removed from the VCF.

#### Selection of reads supporting possible fusions

NanoFG selected candidate SVs that possibly form a fusion gene by annotating both ends of an SV with genes from the ENSEMBL database^24^. If both ends of the SV are positioned in different genes it was flagged as a possible fusion. Next, all the reads supporting the candidate SVs were extracted with samtools (v. 1.7)^25^.

#### Remapping and SV detection

All reads extracted per candidate fusion gene were re-mapped using LAST^26^ with default settings for increased mapping accuracy. Then, NanoSV was used to accurately define the breakpoints in the remapped fusion candidates. NanoSV parameters ‘cluster_count=2, depth_support=False’ were used to detect all present fusions. For C1 and HS-SYII, ‘cluster_count=1’ was used as a parameter for NanoSV.

#### Checking and flagging fusions

Additional information from the ENSEMBL database was gathered to produce an exact composition of the fusion gene. Only fusions that have the ability to produce a continuous transcript on the same strand were retained and additional flags were added to the sample to give extra indication if reported fusions are likely important or if some information from the ENSEMBL database is incomplete.

#### Output generation and visualization

All gathered ENSEMBL gene information was used to produce an overview of the detected fusions. This includes the genes involved, the exon or intron containing the breakpoint, the exact position of the fusion, the number of fusion supporting reads, involved CDS length of both fused genes and the final fused CDS length. The detected fusions were also reported in VCF format for further analysis. To give a better overview of detected fusions, NanoFG also produced a visual overview in PDF format. Apart from information on the genes, flags, position and fusion supporting reads it also included the locations of protein domains to provide quick insight into what domain are involved in the fusion.

#### Primer design

NanoFG automatically designed primers for fusion gene validation using primer3^7^ with default settings, aiming for a 200-400 bp product.

#### Minimal sequencing duration experiment

To detect differences in fusion gene detection based upon sequencing duration, all fastqs were merged and all reads were sorted based on the time of sequencing. The earliest time was taken as the start of the sequencing run and subsequently reads were selected based on bins of 1, 2, 3, 4, 5, 6, 12, 18, 24, 30, 36, 42 and 48 hours after the first read had been sequenced. NanoFG was then run separately on every fastq by using default settings for every sample. Using this approach, the time points where at least 2 supporting reads of a fusion have been sequenced can be determined to define the minimal sequencing duration necessary for each sample to produce two fusion-spanning reads.

## Acknowledgments

We thank all members of the Kloosterman and van Haaften groups for fruitful discussions and support. The authors thank KWF for supporting C.S. and W.P.K. grant UU 2012-5710. This work was supported by funds from the Utrecht University to implement a single-molecule sequencing facility. We thank the Utrecht Sequencing Facility for the Nanopore Sequencing. The colon cancer samples were kindly provided by Prof Ijzermans, Dept of Surgery, Erasmus Medical Center Rotterdam, The Netherlands. Miriam Guillen Navarro, Susan Arentsen-Peters, Heathcliff Dorado-Garcia and Victor Bardinet have kindly helped with providing the clinical samples and information.

## Authors’ Contribution

CS, WPK and GM conceived the study. CS and GM designed experiments, and CS, TV and IR performed the experiments. RCG, AGH and RS provided samples and clinical information for the study. SB, JEV-I and MJR performed bioinformatic analysis. CS and SB analyzed data and CS, GH and GM interpreted the data. CS wrote the manuscript, which was edited by WPK, EEV, GH and GM and reviewed by all authors.

