## Supplemental Figures for "Partner-independent fusion gene detection by multiplexed CRISPR/Cas9 enrichment and long-read Nanopore sequencing"

A

3′ unknown sequence

5′ ATG............AGGATGCTCCATGCTAGCTAGCGGCTAGGCTAGGGACTAATC 3′

3′ TAC............TCCTACGAGGTACGATCGATCGCCGATCCGATCCCTGATTAG 5′

crRNA sequence: 5′TGCTAGCTAGCGGCTAGGCT3′

Read direction: Downstream

B

5′ unknown sequence

5′ ATG............AGGATGCTCCATGCTAGCTAGCGGCTAGGCTAGGGACTAATC 3′

3′ TAC............TCCTACGAGGTACGATCGATCGCCGATCCGATCCCTGATTAG 5′

crRNA sequence: 5′AGCCTAGCCGCTAGCTAGCA3′

Read direction: Upstream

Supplemental Figure 1:

Examples of crRNA design for directional sequencing are shown. **(A)** crRNAs (grey) are designed containing the sequence of the forward strand of the target gene 20 bp prior to a PAM-sequence (NGG, red underlined) if the unknown gene partner is downstream (3’)of the known target sequence. **(B)** crRNAs (grey) are designed containing the sequence of the reverse strand of the target gene 20 bp prior to a PAM-sequence (NGG, red underlined) if the unknown gene partner is upstream (5’) of the known target sequence.

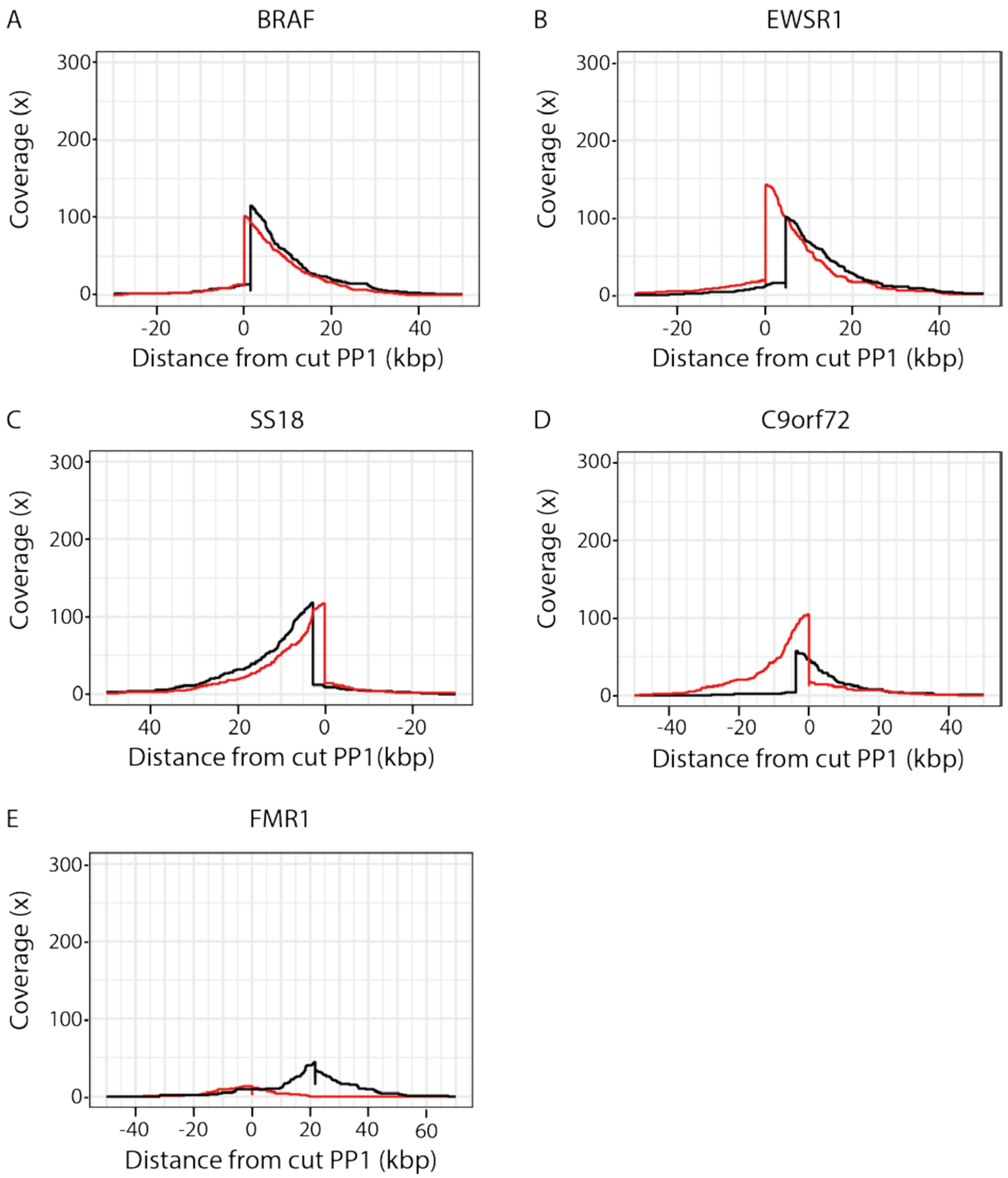


Supplemental Figure 2: **(A-E)** Coverage plots showing deconvoluted on-target coverage across multiple genomic loci for two different cut positions (red = PP1 and black = PP2).

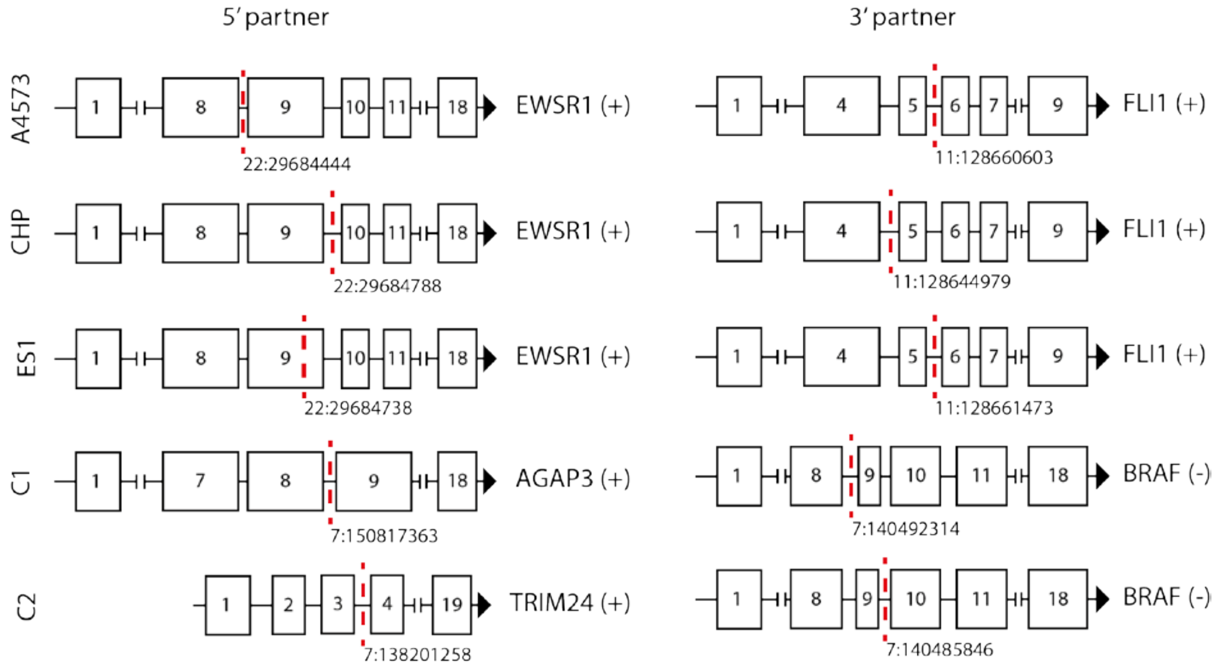


Supplemental Figure 3: Fusion gene configurations for the identified *EWSR1-FLI1*, *AGAP3-BRAF* and *TRIM24-BRAF* fusion genes. (+) or (-) indicate genomic location on the forward or reverse strand of the gene, respectively. Dashed lines (red) indicate break-position for each sample.

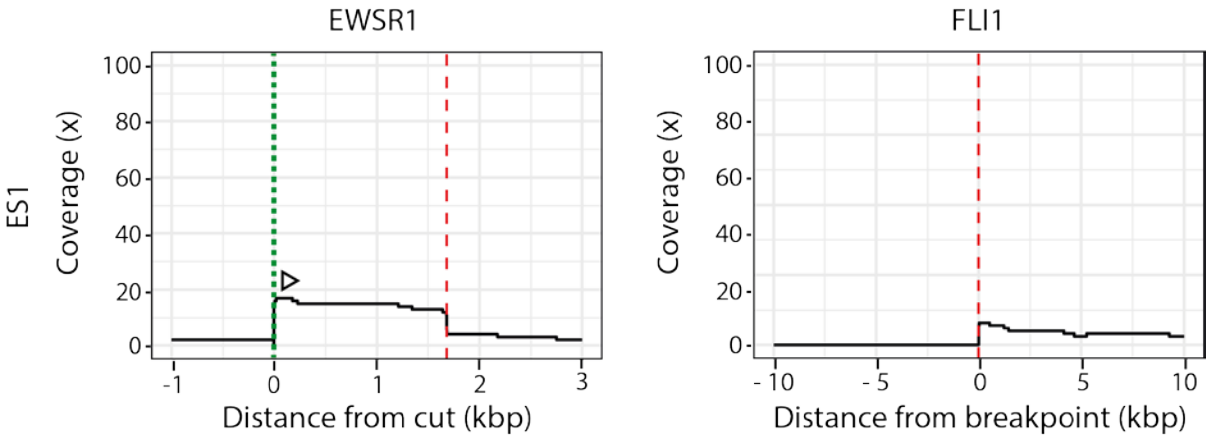


Supplemental Figure 4: Coverage plots for the ES1 tumor sample for the two fusion partners *EWSR1* (targeted) and *FLI1*. Dotted lines (green) indicate cut position, dashed lines (red) indicate breakpoint positions and arrows indicate the desired read direction.

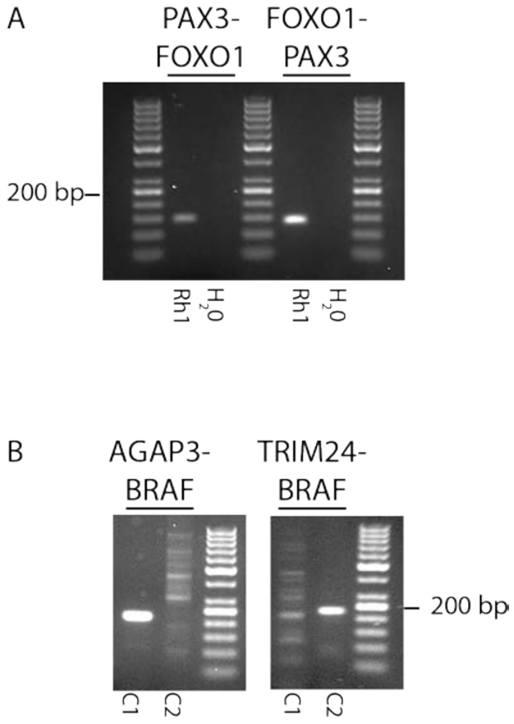


Supplemental Figure 5: Breakpoint PCR for the (**A**) reciprocal *PAX3-FOXO1* and *FOXO1-PAX3* fusion genes in the RH tumor sample and for the (**B**) *AGAP3-BRAF* and *TRIM24-BRAF* fusion genes in the non-amplified tumor material of C1 and C2.

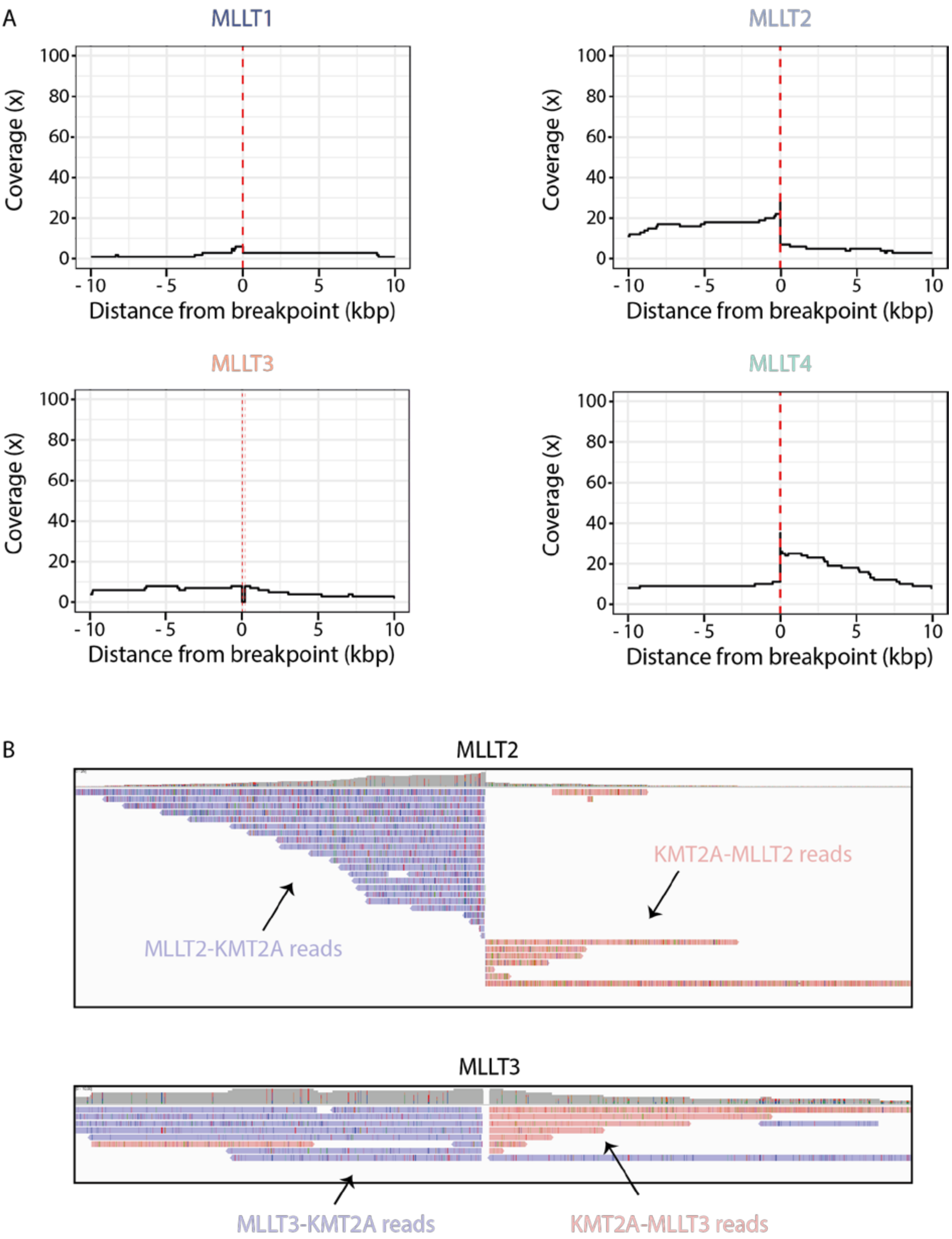


<u>Supplemental Figure 6:</u>

(**A**) Coverage plots for the *KMT2A* fusion partners *MLLT1*, *MLLT2*, *MLLT3* and *MLLT4*. Dashed lines (red) indicate breakpoint positions. (**B**) IGV screenshot showing reads identifying the *KMT2A-MLLT2* and *KMT2A-MLLT3* fusion genes (red) and their reciprocal translocations (blue).

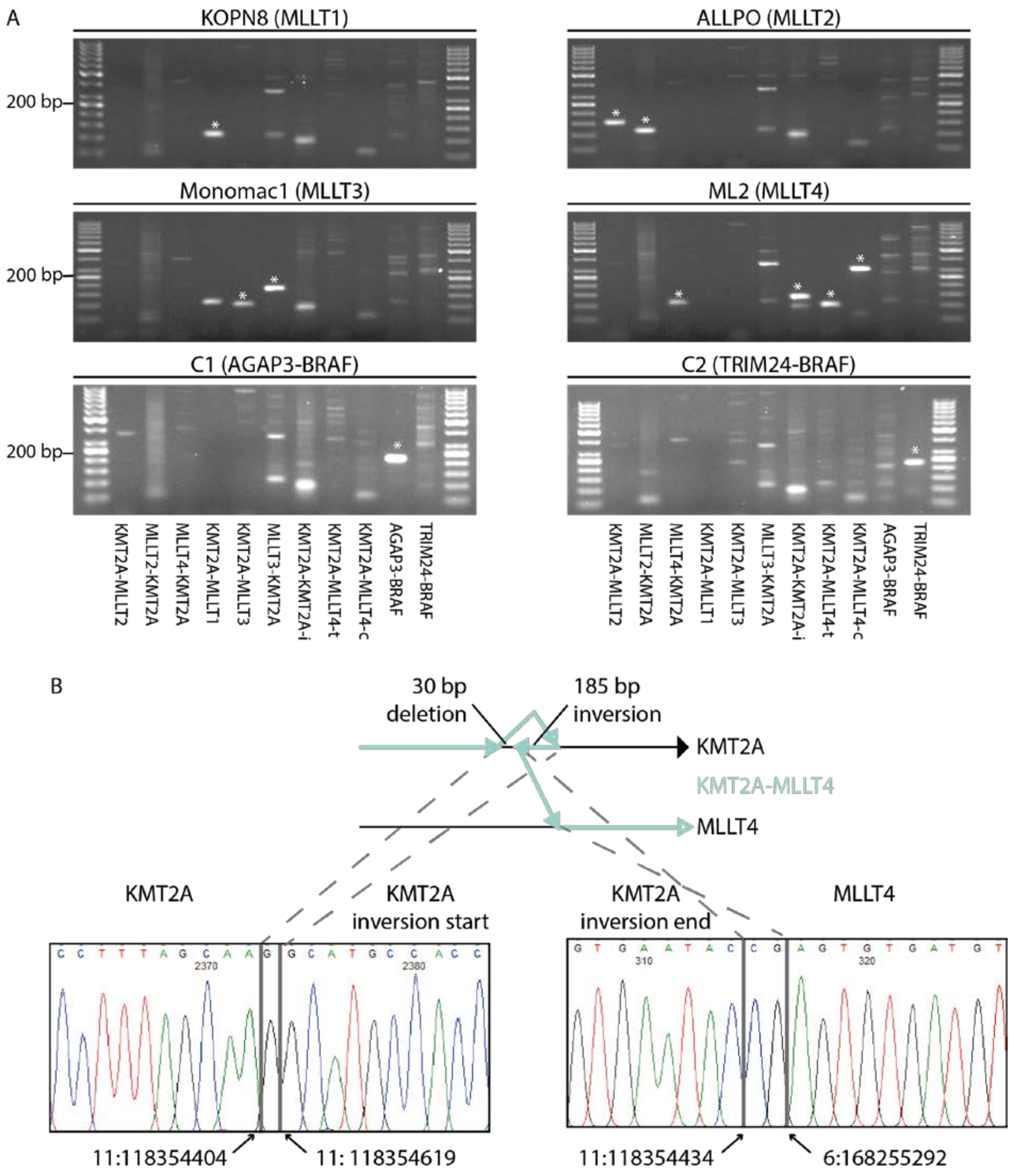


Supplemental Figure 7:

(**A**) Breakpoint PCR with the fusion-specific breakpoint primers for the cell lines KOPN8 (*MLLT1*), ALLPO (*MLLT2*), Monomac-1 (*MLLT3*), ML2 (*MLLT4*) and the tumor samples C1 (*AGAP3-BRAF*) and C2 (*TRIM24-BRAF*). * indicates the bands at the correct height in the correct sample. For the complex *KMT2A-MLLT4* fusion, primers that span the inversion (KMT2A-MLLT4-i), translocation (KMT2A-MLLT4-t) and the complete rearrangement (KMT2A-MLLT4-c) were tested. (**B**) Schematic of rearranged *KMT2A-MLLT4* fusion gene. Sanger-traces showing the exact breakpoints (vertical lines) and breakpoint-positions. The discordant nucleotides between breakpoints (“G” between *KMT2A/KMT2A* inversion; “CG” between *KMT2A* inversion/*MLLT4*) were most likely introduced during non-homologous end joining.
